# Evoked Responses to Localized Sounds Suggest Linear Representation of Elevation in Human Auditory Cortex

**DOI:** 10.1101/2023.05.03.539222

**Authors:** Ole Bialas, Burkhard Maess, Marc Schönwiesner

## Abstract

The auditory system computes the position of a sound along each of the three spatial axes, azimuth, elevation and distance, from very different acoustical cues. The extraction of sound azimuth from binaural cues (differences in arrival time and intensity between the ears) is well understood, as is the representation of these binaural cues in the auditory cortex of different species. Sound elevation is computed from monaural spectral cues arising from direction-dependent filtering of the pinnae, head, and upper body. The cortical representation of these cues in humans is still debated. We have shown that the fMRI blood-oxigen level-dependent activity in small parts of auditory cortex relates monotonically to perceived sound elevation and tracks listeners internal adaptation to new spectral cues. Here we confirm the previously suggested cortical code with a different method that reflects neural activity rather than blood oxigenation (electroencephalography), show that elevation is represented relatively late in the cortex, with related activity peaking at about 400 ms after sound onset, and show that differences in sound elevation can be decoded from the electroencephalogram of listeners, particularely from those who can distinguish elevations well. We used an adaptation design to isolate elevation-specific brain responses from those to other features of the stimuli. These responses gradually increased with decreasing sound elevation, consistent with our previous fMRI findings and population rate code for sound elevation. The long latency as well as the topographical distribution of the elevation-specific brain response indicates the involvement of higher-level cognitive processes not present for binaural cue representation. The differences between brain responses to sounds at different elevations predicted the listeners sound localization accuracy, suggesting that these responses reflect perceived elevation. This is, to our knowledge, the first study that demonstrates the cortical encoding of sound elevation in humans with high-temporal resolution. Our results agree with previous findings from functional magnetic resonance imaging, providing strong support for the hypothesis that elevation is represented in a population-rate code. This represents a critical advance in our understanding of spatial auditory processing along a dimension that is still poorly understood.

## Introduction

The auditory system does not encode the position of a stimulus on the sensory epithelium in the cochlea. Instead, it must observe the movement of the eardrums, caused by the superpositioned sound waves impinging on the ear, and infer source locations from that movement. To this end, the auditory system processes interaural and spectral localization cues and maps them to locations in space. Interaural cues are differences in the sound’s intensity and timing between both ears (Rayleigh, 1907; Sandel et al., 1955). They are computed in the superior olivary complex of the auditory brainstem (McAlpine, Jiang, and Palmer, 2001; Yin, 2002) and determine our perception of a sound’s azimuth. Spectral cues are patterns of peaks and notches in the sound’s spectrum resulting from directional filtering trough head and pinnae (Batteau, 1967; Wightman and Kistler, 1989a; Hofman, Van Riswick, and Van Opstal, 1998). The positions of peaks, notches and edges in the spectrum are computed in the dorsal and posterior ventral cochlear nucleus and indicate the sound’s elevation (Nelken and Young, 1994; Reiss and Young, 2005).

Interaural and spectral cues are integrated in the midbrain where neurons responding to sounds in a narrow region form a topographic representation of auditory space (Middlebrooks and Knudsen, 1984; King and Hutchings, 1987; Gaese and Johnen, 2000). However, most cortical neurons do not respond selectively, but rather modulate their response across a large section of space, preferentially within the contralateral hemifield (Imig, Irons, and Samson, 1990; Middlebrooks et al., 1994; Brugge, Reale, and Hind, 1996; McAlpine, Jiang, and Palmer, 2001). Thus, sound location could be represented in an opponent-channel code where two neural populations are broadly tuned to the contralateral hemifield, encoding location in their relative levels of activity (Salminen et al., 2010; Magezi and Krumbholz, 2010). This model predicts the greatest change in the neural response, and hence highest perceptual resolution, around the midline where tuning curves of both populations intersect. While this may be a plausible mechanism for the perception of azimuth, which is most accurate around the midline, it does not explain the human perception of sound elevation which has a relatively constant accuracy troughout the frontal field (Wightman and Kistler, 1989b; Middlebrooks and Green, 1991).

Instead, elevation could be represented by the overall level of activity within a single neural population. Evidence from fMRI suggests that all elevation-sensitive voxels respond with a similar linear decrease in activity to increasing elevation and that this tuning co-varies with the effects of behavioral manipulations (Trapeau and Schönwiesner, 2018). However because fMRI measures the hemodynamic response which is delayed with respect to neural activity, the latency of cortical elevation processing remains unclear. Also, lying down during scanning might change the neural tuning which is, at least in part, allocentric (Schechtman, Shrem, and Deouell, 2012; Town, Brimijoin, and Bizley, 2017). Thus we used EEG for measuring neural responses with high temporal resolution in a more natural listening situation to test the hypothesis that the cortex represents sound elevation in a linear population rate code.

Unfortunately, EEG, unlike fMRI, can not measure the elevation-sensitive populations in isolation but instead picks up a mixture of all instantaneous neural activity. Our previouse fMRI results suggest that only a fraction of auditory cortex encodes sound elevation (Trapeau and Schönwiesner, 2018). Thus elevation-specific components might be obscured by responses to sound onsets. Indeed, a study that attempted to decode sound location from EEG found that the decoding accuracy for elevation was barely above chance (Bednar, Boland, and Lalor, 2017). To overcome this limitation, we used neural adaptation which is the decay in neural activity following repeated or continuous stimulation due to a variety of physiological mechanisms (Benda, 2021). By playing a longer trail of noise (adapter) before each short stimulus (probe) we cause adaptation of the sound-responsive neurons. Since adapter and probe are cross-faded and identical except for their elevation, the response to the probe is driven by the change in elevation, rather than the over-all sound onset response. We used two different variations of this paradigm, one where the adapter is presented from a loudspeaker and one where it is presented from fully open “hovering"-style headphones. The latter design has the advantage that the adapter is not affected by the directional ear filter and thus does not cause adaptation of elevation-sensitive neurons (Møller et al., 1995).

## Methods & Materials

### Subjects

Twenty-three subjects (15 female) participated in the first and thirty subjects (15 female) in the second experiment. They were between 20 and 33 years old and had no history of neurological or hearing disorders. All subjects gave informed consent and were monetarily compensated at an hourly rate. All procedures were approved by the ethics committee of the medical faculty at the University of Leipzig (reference number 248/17-ek).

### Apparatus

Subjects sat in the center of a custom-built spherical array of loudspeakers (model Mod1, Sherman Oaks, CA, USA) inside a 40 m^2^ hemi-anechoic chamber (Industrial Acoustics Company, Niederkrüchten, Germany). Two additional speakers were mounted hovering next to the subject’s ears, pointed at the ear canal, and used as fully open headphones in the second experiment. Because of their proximity and orientation, sounds from those loudspeakers were unaffected by the listeners directional transfer function (Møller et al., 1995). This allowed us to simultaneously present sounds from loudspeakers at different elevations in the array as well as non-spatial headphone sounds. We equalized the transfer functions of each loudspeaker by applying an inverse filter to the stimuli upon presentation. Two digital signal processors and six 8-channel amplifiers (models RX8.1 and SA8, Tucker-Davis Technologies, Alachua, FL, USA) drove the loudspeakers at 50 kHz sampling rate. The processors’ digital ports controlled the LEDs attached to each loudspeaker, obtained responses from a custom-built button box, and sent event triggers to the EEG. We used a 64-channel system (model BrainAmpMR, Brain Products, Gilching Germany) to record EEG at a sampling rate of 500 Hz. The active silver/silver chloride electrodes were fixed on the subject’s head with an elastic cap (Easycap, Germany) according to the international 10/20 system with FCz as reference electrode. Electrode impedance was kept below 2 kΩ. Two cameras (model Firefly S, Teledyne FLIR, OR, USA), positioned between loudspeakers, were used to monitor the subjects’ head pose.

### Software

We programmed the digital signal processors using Real-time Processor Visual Design Studio (Tucker-Davis-Technologies, Alachua, FL, USA) and controlled the cameras using the Spinnaker SDK (Teledyne FLIR, OR, USA). The software of both devices provides an API that we integrated in a custom Python module that controlled the experimental apparatus (Bialas, 2022). We estimated subjects’ head pose by capturing an image of their head, localizing facial landmarks with a deep neural network, implemented in Pytorch (Paszke et al., 2017), and mapping these points to a generic 3D model using functions from the OpenCV library (Bradski, 2000). Stimuli and trial-sequences were generated using the slab Python module for psychoacoustics (Schönwiesner and Bialas, 2021). We recorded EEG signals using Brain Vision Recorder (Brain Products, Gliching Germany), and imported them into MNE-Python for analysis (Gramfort et al., 2013). A full list of the software environment can be found in the accompanying [online repository](https://github.com/OleBialas/elevation-encoding).

### Preprocessing

A de-noising algorithm, combining filtering and source separation, removed power line artifacts while minimizing temporal distortions due to filtering (Cheveigné, 2020). We then applied a causal, minimumphase, high-pass filter with a hamming window and a 1 Hz cutoff frequency. After epoching (without applying a baseline), channels in which the correlation between the actual signal and the signal predicted by the neighboring channels was less than 0.75 for more than 40 percent of the time (Bigdely-Shamlo et al., 2015) were replaced with interpolated data from the surrounding channels using spherical splines. We then subtracted the average signal across all channels from each channel (“average reference”). Next, independent component analysis unmixed the signal, and an algorithm removed components corresponding to eye-blinks based on their topographical correlation with a previously determined template (Viola et al., 2009; Plöchl, Ossandón, and König, 2012). Finally, an algorithm determined channel-specific rejection thresholds and repaired or removed epochs where the thresholds were exceeded (Jas et al., 2017). Except for the initial selection of a reference for blink removal, the entire preprocessing pipeline was automated. Afterwards, the data were inspected by eye to assess the effect of preprocessing.

### Experiment I

Subjects sat comfortably on a height-adjustable chair in an anechoic chamber. Target loudspeakers stood at a distance of 3.2 m at elevations of 37.5°, 12.5°, −12.5° and −37.5° with respect to the subject’s interaural plane. Because perception of sound source elevation is slightly more accurate for lateral targets (Makous and Middlebrooks, 1990), all target speakers were positioned at an azimuth of 10° to the subject’s right. Initially, we tested subjects’ ability to localize sounds and familiarized them with the setup. To avoid that subjects explicitly learn the target speakers’ directional transfer functions during training, we used a different set of loudspeakers located at elevations of 50°, 25°, 0°, −25° and −50°. In each of the 200 test trials, subjects heard a 150 ms burst of noise with a 5 ms on- and offset ramp from one speaker. Subjects localized each sound by pressing one of four buttons on a custom-built box. There was no time limit for responding, and the subsequent trial started automatically after the subject had responded to the previous stimulus. The order of speakers was randomized without direct repetition of the same speaker. Subjects were instructed to keep their head and gaze aligned with the fixation cross at 0° azimuth and elevation. After completion of the test, we prepared the EEG electrodes. During recordings, each trial started with 600 ms of noise (adapter) played from the speakers at either 37.5° or −37.5°. Then, a 150 ms burst of noise played from one of the other speakers, resulting in six different adapter-probe pairs. Adapter and probe had overlapping 5 ms on- and offset ramps so that the sound intensity remained constant during the transition. The adapter’s initial position was chosen randomly and changed every 30 trials. The probe’s location was chosen randomly without direct repetitions of the same speaker. Every probe was followed by a 350 ms silent inter-stimulus interval. In five percent of all trials, the probe did not come from one of the target positions but from a random speaker within the frontal field. Subjects had to respond to these deviant trials by pressing a random button as fast as possible. If they managed to respond within one second after sound onset, the trial was considered a success. After one second, the trial was considered failed, and stimulation resumed. The experiment was divided into four blocks, each of which consisted of 504 trials and lasted 35 min in total. We instructed subjects to keep their head and gaze aligned with the fixation cross throughout the recording but we did not check whether they complied.

### Experiment II

Again, subjects completed an initial test in which they had to localize sounds coming from loud-speakers at 50°, 25°, 0°, −25° and −50°. In each of the 30 trials, subjects heard a 100 ms from one of the speakers which they localized by pointing their head in the direction of the speaker and pressing a button. This triggered the cameras to acquire images from which the head-pose was estimated. After localizing the sound, subjects had to return to the central fixation cross and press the button again to start the next trial. If their head pose was not aligned with the fixation cross a warning tone prompted them to adjust their position. The first 15 stimuli were accompanied by a visual cue (i.e. a flashing LED at the loudspeakers’ location) and familiarized subjects with the procedure. The stimulus protocol during recording was similar to the experiment I except for the duration of adapter (1000 ms), probe (100 ms) and inter-stimulus interval (900 ms). Also, the adapter was played from headphones instead of one of the target speakers. In five percent of trials, subjects heard a tone after the inter-stimulus interval which prompted them to localize the probe they had just heard. Thus, they had to pay attention to each probe’s location since they did not know whether they would be asked to localize any given sound. The experiment was divided into six blocks, each consisting of 240 trials, roughly lasting 55 min in total.

### Elevation Gain

To quantify the accuracy of the subjects’ perception of elevation, we used the elevation gain (EG) which is the slope of the regression between the target and response elevation. Thus, the EG measures how strongly subjects modulate their response with changes in the target’s elevation which 1 indicating perfect localization and 0 indicating random responses. We chose EG because it robust to outliers and has been shown to capture changes in behavior and neural tuning following manipulation(Hofman, Van Riswick, and Van Opstal, 1998; Trapeau and Schönwiesner, 2018). We computed the EG for the localization tests in both experiments as well the task in experiment II. In the latter, the EG for two subjects, which was slightly negative, was set to zero for subsequent analyses.

### Permutation Testing

When investigating a phenomenon with unknown latency and locus it is desirable to test for differences between conditions across all points in time and space while controlling the false alarm rate. This is achieved by a permutation-based cluster test which operates under the null hypothesis that all conditions are exchangeable and finds spatio-temporal clusters that violate this assumption (Maris and Oostenveld, 2007). Unsing this algorithm, we calculated F-scores, selected samples where the F-score exceeded a threshold corresponding to an uncorrected p-value of 0.05 and clustered them for temporal and spatial adjacency. The size of an observed cluster is defined by the sum of all F-scores. By repeating this procedure on randomly permuted data, we obtain a distribution of cluster sizes that occur by chance. The p-value of the actually observed effects is given by the probability of observing a similar or larger effect under the permutation distribution. Thus, the cluster test is sensitive while avoiding the multiple comparisons problem because statistical inference is constrained to the distribution of cluster sizes. This however means that the spatial and temporal extent of the clusters is purely descriptive and not statistically controlled (Sassenhagen and Draschkow, 2019). We used the cluster test implemented in MNE-Python with 10000 permutations. We conducted one test per subject, comparing all conditions simultaneously. If the test returned more than one significant cluster for a subject we ignored all but the largest.

### Decoding

While significant clusters indicate differences between conditions, they do not indicate which conditions differ in what way. Thus, to complement permutation testing, we used multivariate logistic regression to decode sound elevation from brain responses. We chose logistic regression because it is simple, well-understood, and robust to overfitting (Dreiseitl and Ohno-Machado, 2002). The multivariate logistic function was fit to the data sample-by-sample using ridge-regression, and denotes the probability of a given observation (i.e. the instantaneous distribution of voltage across channels) belonging to either one of two classes (i.e. elevations). To avoid picking an arbitrary decision threshold, we used the receiver operating characteristic (ROC) curve, which denotes the ratio between true and false positives across all possible thresholds. Thus, the area under the ROC curve is an unbiased measure of accuracy with 1 indicating perfect and 0.5 indicating random classification. To avoid overfitting we used 100-fold leave-one-out cross-validation, splitting the data into 100 segments, fitting the logistic function to 99 of them and testing it on the left out segment. Each segment was used for testing once, and the accuracy was averaged across all segments. Finally, we used the bootstrap method to resample the subject-specific decoding accuracies 10000 times (Efron, 1992). The resampled data’s mean is an estimate of the group’s average decoding accuracy across time, and its standard deviation indicates the uncertainty of the mean estimate. We used the logistic regression implementation from the scikit-learn package (Pedregosa et al., 2011) and the sliding estimator function from MNE-Python, which applied the classifier sample-by-sample.

### Elevation Tuning

Our main objective was to estimate how brain responses to sounds change with elevation. To this end, we selected the channels where the difference between conditions (i.e. the F-score) was largest in the time interval after subjects heard the probe. Then, we estimated the relationship between average ERP amplitude and sound elevation for all subjects using ordinary least squares linear regression. To quantify the uncertainty in the estimated relationship we resampled the data 10000 times using the bootstrap method, applied linear regression to each resampled set, and computed the standard deviation. Since the bootstrapped sampling distribution is normal, over 95 percent of all estimated linear models lie within two standard deviations of the mean. However, since EEG-sensors measure Voltage relative to the reference, an observed change at any channel could also reflect a change in the reference (i.e. the average of all channels) of opposite sign. To overcome this limitation we computed the current source density (CSD) using the implementation from MNE-Python, which is based on spherical spline surface Laplacians. The CSD is a reference-free estimate of radial current flow at the scalp where positive values represent outward (i.e. from the brain to the scalp) and negative values inward current flow (Kayser and Tenke, 2015). Additionally, it acts as a spatial high-pass filter suppressing the effects of volume conductance and improving the spatial resolution (Cohen, 2014). This effectively simplifies the variance structure of the data making CSD a useful preprocessing step for principle component analysis (Kayser and Tenke, 2006). PCA transforms the data to a n-dimensional orthogonal space so that each dimension accounts for the largest possible share of variance. Thus, the principle components are patterns of covariance reflective of underlying neural generators that generated the observed data (Cohen, 2014). This follows the rational that channels co-vary because they area affected by the same generator which might be a localized source or a distributed network. PCA is a simple and well-understood method for data-driven detection of relevant components and has been shown to produce meaningful summaries of average evoked responses (Donchin, 1966; Chapman and McCrary, 1995; Kayser and Tenke, 2006).

### Lowest sounds cause the largest responses

Participants were able to localize sounds accurately as shown by an average EG of 0.74(*SD* = 0.19) in the initial localization tests, which is similar to what was reported in previous studies (Hofman, Van Riswick, and Van Opstal, 1998; Trapeau and Schönwiesner, 2018). During the continuous sound presentation in experiment 2, uncertainty about target trials and added memory load reduced the average elevation gain significantly compared to the performance in the localization test just before (0.78 and 0.45, (*t*(29) = 6.05, *p* < 0.001).

Neural responses generally increased with decreasing sound elevation in both experiments although the effects differed in size and latency. In the first experiment, elevation-specific event-related potentials (ERPs) differed mostly in the time interval from 100 ms to 300 ms after probe onset (colored bar in 1A). This was confirmed by a permutation-based cluster test, which compared the responses to all six adapter-probe combinations (adapter at 37.5° and probes at 12.5°, −12.5°, −37.5° and adapter at −37.5° and probes at −12.5°, 12.5°, 37.5°). The test found significant clusters, comprising most of the electrodes, for 9 of the 23 subjects. Examination of the F-scores, averaged across time between 100 ms and 300 ms (Fig.1B) revealed that the difference between elevation-specific ERPs was largest at the central electrode Cz. Thus, we selected this electrode to investigate how the ERP changed with elevation more closely.

**Figure 1.**
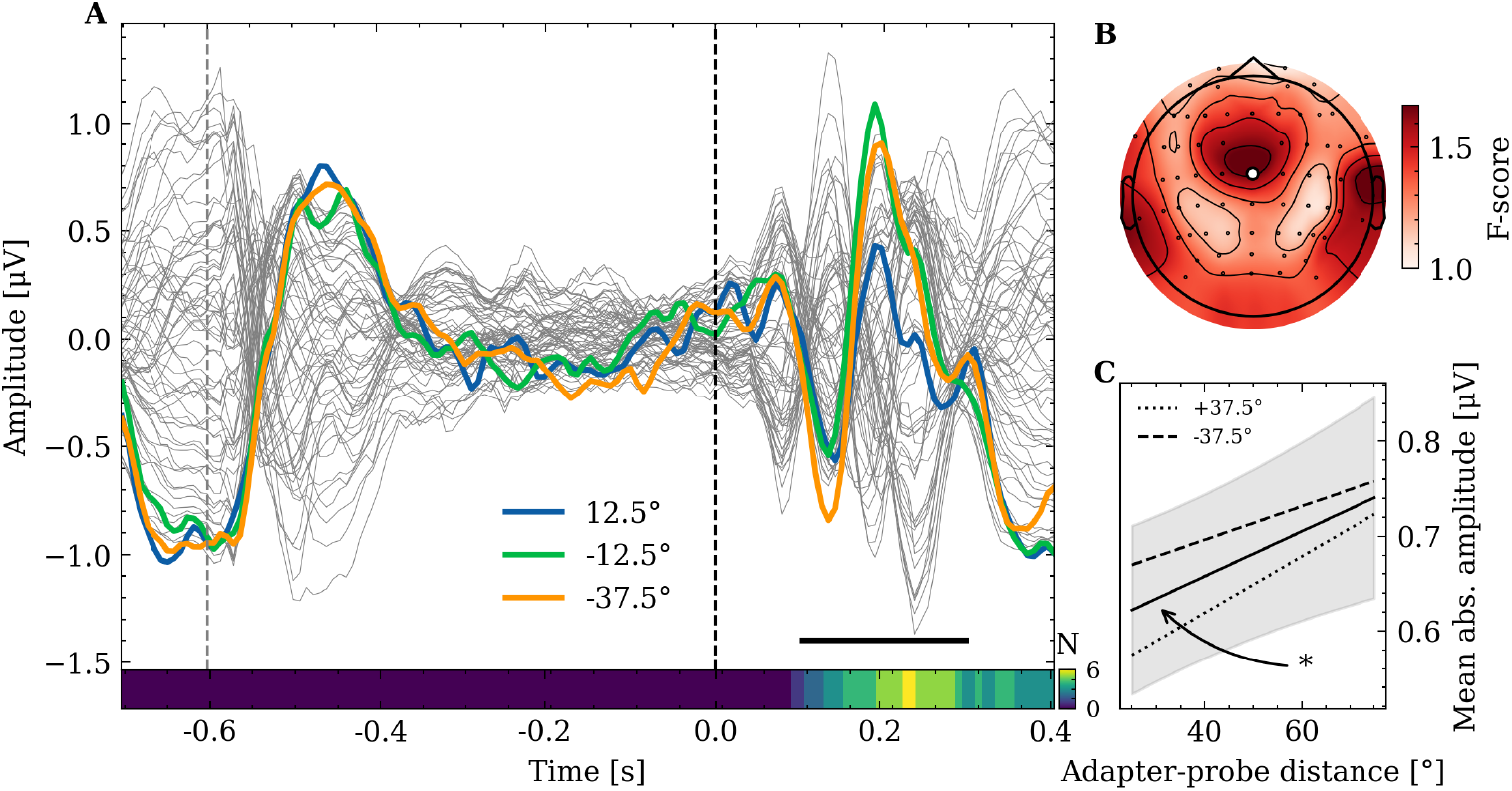
**A**: Grand average evoked response from the first experiment. Gray lines show the voltage at each channel averaged across sound sources and colored lines show the response to each probe at the channel Cz where the difference between conditions was largest (white dot in **B**), when the adapter was played from 37.5°. The gray dashed line marks the onset of the adapter and the black dashed line the transition to the probe. The horizontal color bar indicates the number of subjects for whom the cluster-test showed a significant differences between conditions at any point in time. **B**: topographical distribution of F-scores. Color represents the average F-score at each channel in the time interval marked by the horizontal black line in **A**. The white dot indicates the channel with the largest score (Cz). **C**: relationship between adapter-probe separation and average absolute ERP amplitude. The solid gray line shows the change in average absolute ampltidue across all sound sources and the dotted and dahes lines show the change across all sources when the adapter located at −37.5° and 37.5° respectively. The shaded interval shows ±2*SD*, estimated via bootstrapping.

The average ERP at Cz showed two deflections, of opposite polarity, which increased in amplitude with the separation of adapter and probe (Fig.1A). To quantify this trend, we computed the average absolute ERP amplitude in the time interval between 100 ms and 300 ms after probe onset for each adapter-probe combination and regressed it against the distance between adapter and probe (Fig.2. The average absolute amplitude significantly increased with separation of the adapter and probe (*R* = 0.18, *p* = 0.033). This increase was steeper when the adapter was located at 37.5° (*β* = 29.8 * 10−4) compared to when the adapter was located at −37.5° (*β* = 17.7 * 10−4). Even though the trend was not significant when considering only those trials where the adapter was played at 37.5° or −37.5° (*R* = 0.22, *p* = 0.06 and *R* = 0.14, *p* = 0.26), this observation is compatible with a monotonic population-rate code which predicts that elevation-sensitive neurons respond maximally to the lowest sound (i.e. the adapter at −37.5°), rendering them insensitive to subsequent sounds from higher elevations.

**Figure 2.**
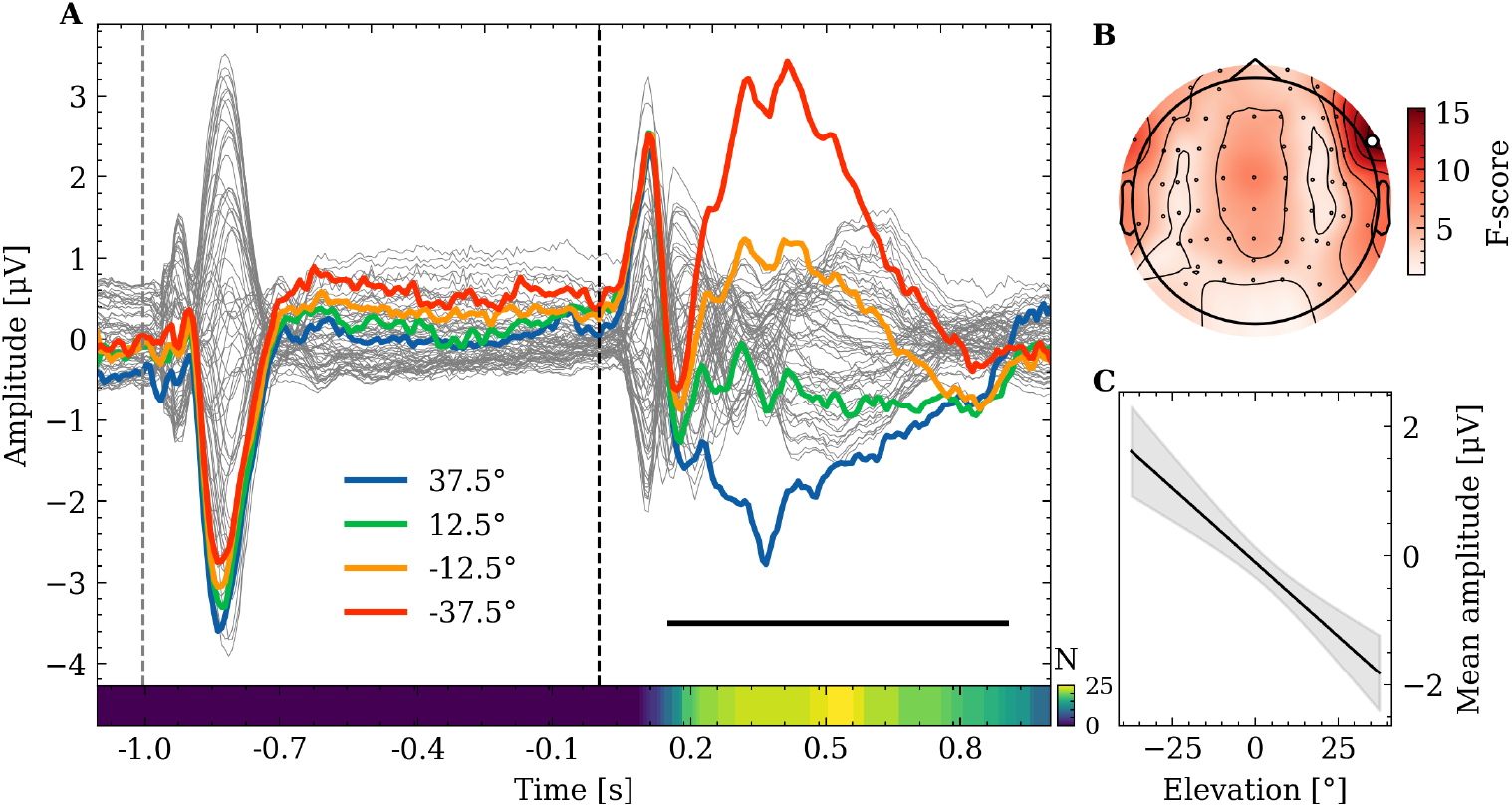
**A**: Grand average evoked response from the second experiment. Gray lines show the voltage at each channel averaged across conditions and colored lines show the response to each probe at the channel FT10 where the difference between conditions was largest (white dot in **B**). The gray dashed line marks the onset of the adapter and the black dashed line the transition to the probe. The horizontal color bar indicates the number of subjects for whom the cluster-test showed a significant differences between conditions at any point in time. **B**: topographical distribution of F-scores. Color represents the average F-score at each channel in the time interval marked by the horizontal black line in **A**. The white dot indicates the channel with the largest score (FT10). **C**: relationship between probe elevation and average ERP amplitude. The shaded interval shows ±2*SD*, estimated via bootstrapping.

The second experiment addressed the confound between adapter and probe position by using a non-spatial adapter. Consequently, the results offered clearer support for the population-rate code hypothesis. The permutation test which compared responses to the different probes at 37.5°, 12.5°, −12.5° and −37.5° elevation found significant clusters for most subjects between 200 ms and 800 ms with a peak around 500 ms after probe onset (colored bar in Fig.2A). The scalp-distribution of F-scores averaged between 150 ms and 900 ms after probe onset (Fig.2B) revealed that the difference between elevation-specific ERPs was largest at fronto-temporal electrodes with a peak at FT10. Again, we chose this electrode for a closer inspection of elevation-specific changes in the ERP.

The average ERP at FT10 exhibited a sustained deflection that was strongly modulated in amplitude by the probe’s elevation (colored lines in Fig.2A). The amplitude was largest when the probe was located at −37.5° and smallest when it was located at 37.5 Linear regression found that the relationship between the probe’s elevation and the average response amplitude in the time interval from 150 ms to 900 ms after onset was highly significant (*R* = −0.66, *p* < 0.001).

### Elevation decoding accuracy predicts task performance

Sounds at different elevations evoked visibly different ERPs. Thus, a logistic regression classifier which decoded sound elevation from EEG performed above chance for all sound source pairs (Fig.3A). Decoding accuracy increased with distance between adapter and probe and was not dependent on the sounds’ absolute elevation (e.g. the curves for 37.5° vs 12.5° and −37.5° vs. −12.5° were virtually identical). Decoding accuracy followed a similar time course in all conditions: it remained at chance level while the adapter was presented, started to increase around 200 ms and peaked around 400 ms after probe onset. Participants who were better at distinguishing sounds at different elevations also had more distinctive ERPs in response to sounds from these different elevations, allowing better decoding accuracy. Subjects’ average decoding accuracy between 150 ms and 900 ms after probe onset correlated with their elevation gain during the experimental task (*R* = 0.51, *p* = 0.004). Notably, this relationship is mostly due to the upper right quadrant in Figure 3B being empty, meaning there were no subjects who’s brain responses were decodable but who failed at the task. There were, however, several subjects for whom decoding failed but who still performed the task accurately.

**Figure 3.**
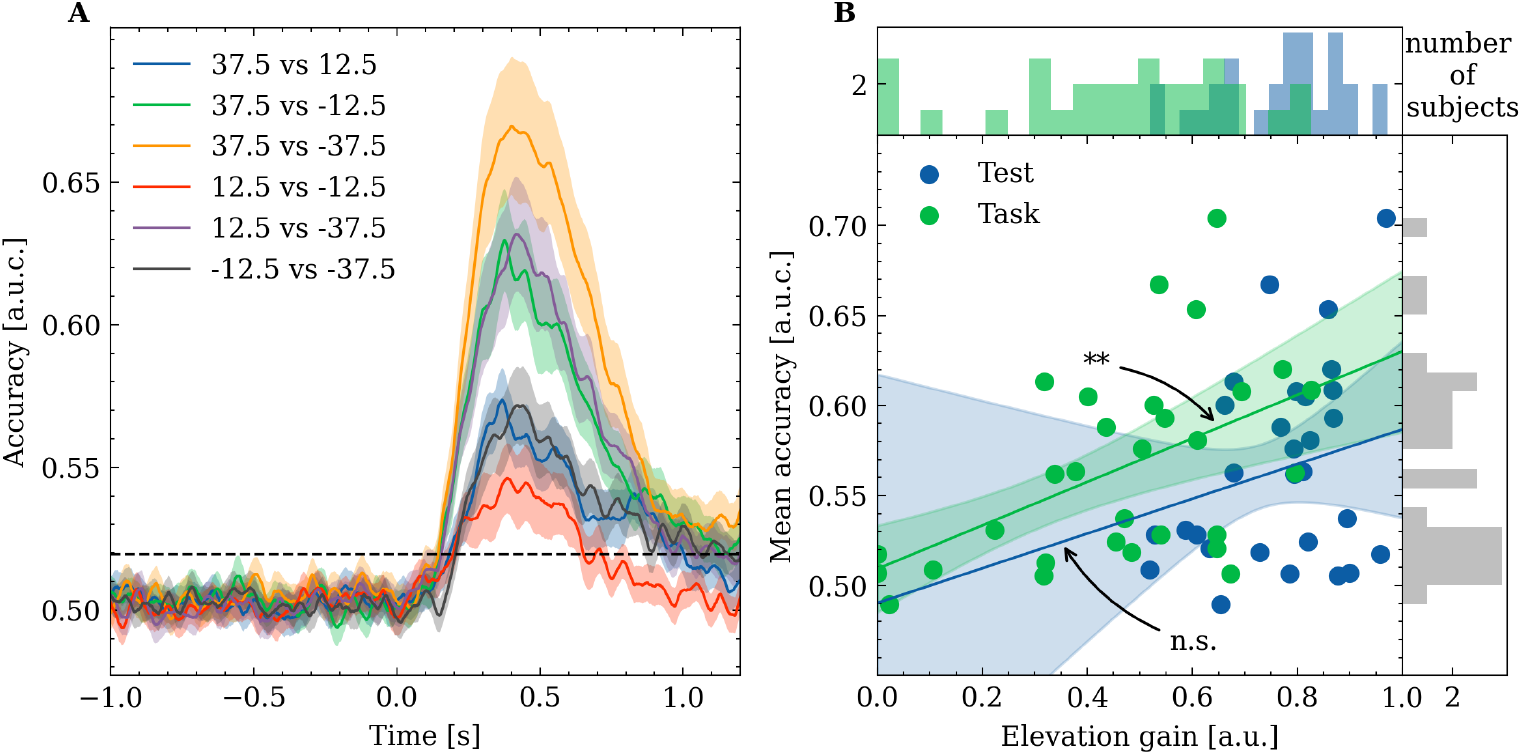
**A**: accuracy across time for decoding the ERPs to all possible pairs of probes. The displayed data are averages across all cross-validation splits. **B**: relationship between decoding accuracy and localization behavior. The lines show the average relationship between decoding accuracy and EG during the localization test (blue) and experimental task (green). Shaded intervals show ±2*SD* estimated via bootstrapping. Bar plots on the right show the marginal distributions of EG during the test (blue), EG during the task (green) and average decoding accuracy (gray).

We also regressed the average decoding accuracy against the EG during the initial localization test to investigate whether the relationship between decoding and performance was specific to the experimental task or applied to sound localization in general. While linear regression revealed a positive relationship between decoding accuracy and localization test EG, this trend was not significant (*R* = 0.23, *p* = 0.23), likely due to the small number of trials and lack of variance across subjects in the localization test.

### Auditory component activates with decreasing elevation

Because ERP morphology depends on the chosen reference it is unclear whether the observed changes in ERP amplitude with elevation reflect a decrease or a change in polarity with respect to the underlying current flow. To resolve this ambiguity we computed the current source density (CSD), a reference-free estimate of current flow, and performed a principle component analysis. The principle component accounting for most variance had a topography suggesting an auditory origin and showed a deflection which gradually increased in amplitude for decreasing elevation (Fig.4A. The second and third components might reflect activity of the motor and prefrontal cortices respectively (Fig.4B&C). These additional components were unaffected by elevation. Together, the three components accounted for 88 % of the variance in the average evoked response.

**Figure 4.**
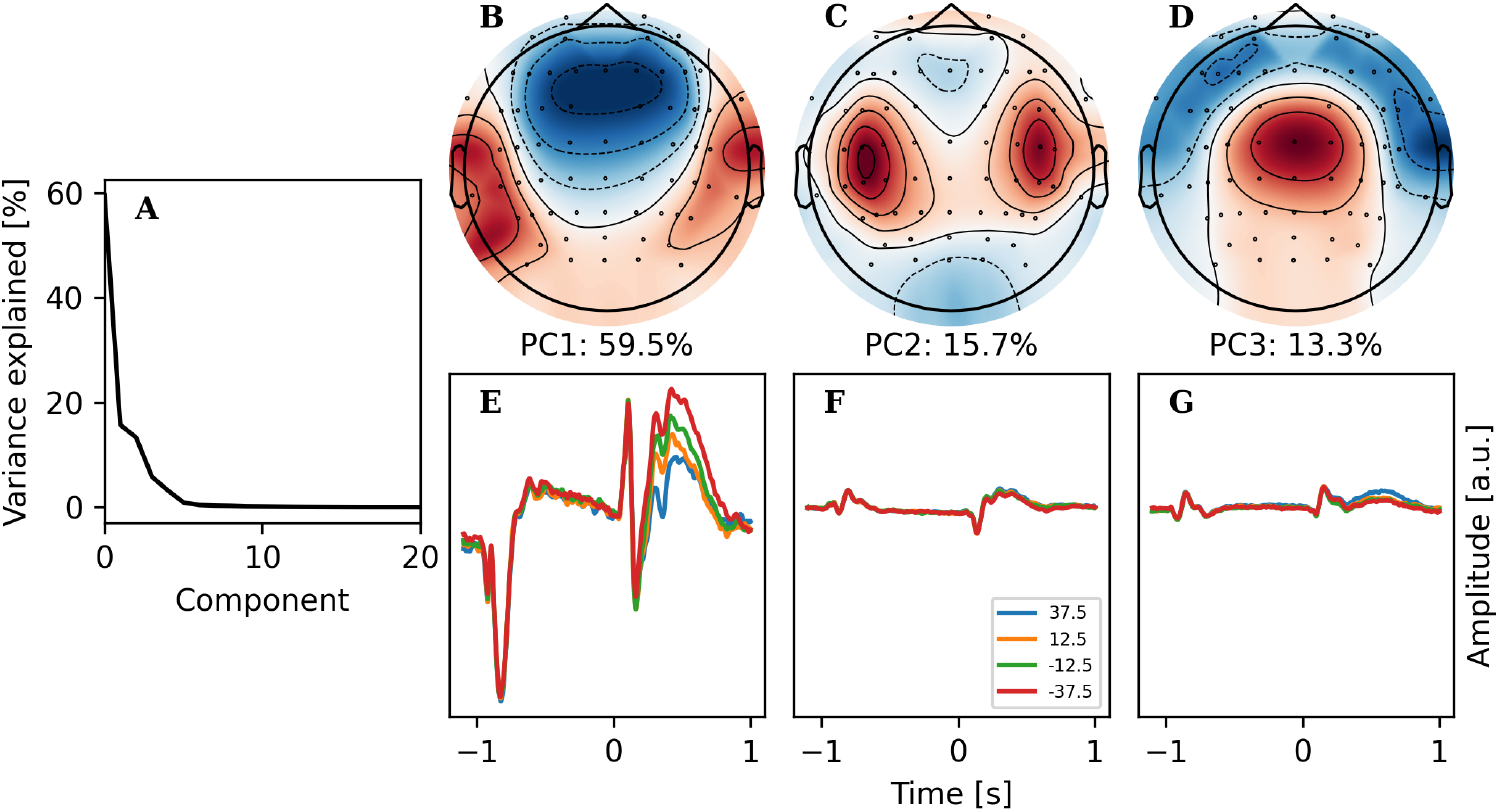
Principle components of the average current source density. **A**: the percentage of variance accounted for by each component. **B-D**:Channel weights of the components as topographical distribution on the scalp. The percentage indicates the total variance accounted for by the respective component. **E-F** Component loading across time for each sound elevation.

## Discussion

### Evoked responses encode sound elevation

Both experiments demonstrate that the cortical processing of sound elevation can be assessed with EEG. Using fMRI, we previously identified voxels in auditory cortex tuned to sound elevation (Trapeau and Schönwiesner, 2018), but the lower spatial resolution of EEG would not have allowed us to isolate the responses from these small patches of auditory cortex. Indeed, a previous EEG study attempted to decode sound elevation from EEG responses and found that decoding accuracy exceeded chance level just barely for only some of the listeners (Bednar, Boland, and Lalor, 2017). The adaptation design helped to circumvent this issue by separating neural activity related to sound onset and elevation in time. In the second experiment, we showed that changes in sound elevation evoke distinct cortical responses that can be decoded accurately, and that decoding accuracy predicts individual localization performance. Decoding accuracy for all pair-wise elevation comparisons followed the same time course with a single peak at around 400 ms after the elevation information became available. The time course and the fact that accuracy was predictive of task performance suggests that we decoded a physiological process distinctly related to elevation rather than processes associated with acoustical features that co-vary but are not causally related with sound elevation. Elevation-specific activity was well captured by a single principle component, suggesting that the observed encoding of elevation reflects a distinct physiological process.

### A monotonic population-rate code for elevation

We hypothesized that the auditory cortex represents sound elevation in a population response that decreases monotonically with increasing elevation. Both experiments presented here offer support for this hypothesis. In experiment 1, ERP amplitude increased with separation of the adapter and probe when the adapter was played from the highest position. Under the assumption of a rate code with minimal response at high elevations, this adapter would evoke small responses and enable increasingly larger responses for probes at lower elevations, with corresponds to the observed pattern. Consequently, the adapter played from the lowest elevation would lead to strong adaptation independent of probe elevation. Thus, as expected, differences between elevation-specific ERPs were larger when the adapter was located at a high elevation. Unexpectedly however, probes tended to evoke larger responses when the adapter was located at a low elevation. This may be explained by co-varying acoustic factors contributing to the probe response and it does not contradict the representation we propose since absolute amplitude does not carry information in a rate code. To summarize, while experiment 1 showed that elevation is reflected in the ERP, its interpretation is limited by the fact that responses are affected by both the adapter and the probe’s position.

The purpose of experiment 2 was to remove the interaction of adapter and probe position by using a non-spatial adapter (see Methods). Consequently, elevation-specific modulation of ERP amplitude was about an order of magnitude larger compared to the first experiment, and this method allowed direct measurement of monotonic elevation tuning functions. Sounds from the lowest elevations evoked the largest response, and response amplitude gradually declined with increasing elevation in a way that was well described by a linear model. The sign of the elevation-specific response amplitude changed around the listener’s eye level. Because EEG-electrodes measure voltage relative to a common reference, it is unclear whether this reflects a gradual decrease or change in polarity of the underlying neural current. We were able to decide between these alternatives by computing a reference-free estimates of neural current flow (current source density), which benefits topographical localization (Kayser and Tenke, 2015). One principle component accounted for more than half of the ERP variance and showed a topography that suggested it originated in the auditory cortex. This component showed a gradual increase in response amplitude with decreasing elevation, supporting the idea of a monotonic population rate code. Two additional components, which together with the first accounted for almost 90 % of the variance, showed topographies suggesting they originated in motor and frontal cortices. Involvement of those regions plausible given that the experimental task required the participants to translate their auditory perception into goal-directed head movements.

Thus, our results bear striking similarity to changes in the BOLD-response to elevated sounds (Trapeau and Schönwiesner, 2018). This is remarkable given the fact that many neural phenomena are not similarly reflected in EEG and fMRI. For example, increases in BOLD-activity may not be picked up by EEG if the sources are not synchronized or their geometric orientation prevents a summation of their local field potentials (Buzsáki, Anastassiou, and Koch, 2012), and increases in EEG power can reflect an overall decrease of the underlying neural activity (Musall et al., 2014). Thus, our combined findings constrain the cortical encoding of sound elevation to sources that affect EEG and fMRI signals in the same way. They also suggest that the cortical encoding of elevation is egocentric, because fMRI recordings in supine and EEG recordings in upright position revealed a similar trend.

### Latency of cortical elevation processing

In the first experiment, we found the largest elevation-specific response between 200 ms and 250 ms after probe onset. This latency is similar to other reports on the cortical processing of elevation. Using MEG, Fujiki and colleagues found that unexpected sounds, deviating in elevation, caused mismatch responses between 150 ms and 250 ms (Fujiki et al., 2002). Bednar and colleagues found that some participants’ brain responses to two differently elevated sound sources could be decoded above chance between 200 ms and 400 ms after sound onset (Bednar, Boland, and Lalor, 2017). Notably both those studies also reported correlates of sound azimuth which occurred earlier in time reflecting the fact that binaural cues depend on precise temporal information while spectral cues result from a detailed analysis of spectral patterns.

In the second experiment, the elevation-specific component started at a similar time but lasted much longer, so that most subjects showed significant differences across ERPs between 200 ms and 800 ms. This is reflected in the time-course of decoding accuracy, which peaked around 400 ms after probe onset. Remarkably, elevation-specific ERPs could still be distinguished 1 s after probe onset, which means that the brain could access the relevant perceptual representation at the end of a trial when participants were informed whether they had to indicate the sound location in this trial. Thus, the increased duration of elevation-specific responses might be a result of the experimental task where maintaining the probe’s perception facilitated performance. Interestingly, visual research identified late ERP components that scale with task-difficulty in latency and magnitude in a way that is predictive of individual performance (Philiastides and Sajda, 2006; Philiastides and Sajda, 2007). Such perceptual persistence could be implemented by feedback loops which reverberate the neural representation after stimulus offset (VanRullen and Koch, 2003; Large, Aldcroft, and Vilis, 2005). The perceptual persistence of a visual stimulus is inversely related to its duration and intensity (Coltheart, 1980). Thus, perceptual persistence could provide a mechanism of evidence accumulation for perceptual decision making under difficult conditions. However, the present study was not designed to investigate the effects of task-difficulty and further research is required to answer those questions.

### The cortical representation of sound direction

It is thought that the auditory cortex represents sound azimuth as the difference between the rates of activity in two opponent neural channels, each tuned to the contralateral hemifield so that most azimuth sensitive neurons respond maximally to cues outside the range created by the head (McAlpine, Jiang, and Palmer, 2001). This may sound counter-intuitive but, in a rate code, accuracy is not related to the absolute magnitude but rather the change in neural response. Thus, placing the tuning curve’s peak outside the physiological range places the slope in the center, where azimuthal localization is most accurate (Harper and McAlpine, 2004). Similarly, the fact that sounds at low elevations cause larger responses than sounds at high elevations does not mean that they are localized more accurately. Instead, the steady slope of the population response suggests that accuracy remains constant across elevations, which lines up well with localization behavior (Wightman and Kistler, 1989b; Middlebrooks and Green, 1991). There is no cardinal reason why the maximum of the tuning is at low rather than high elevations. While it is tempting to argue that this may be due to a tendency for deeper spectral notches at low elevations, this explanation was disproved by our previous finding that the tuning function depends on perception rather than the acoustic nature of the cues: the tuning curve flattens when listeners are presented with spectral cues from other ears, and they re-emerge as the listener adapts to the new cues (Trapeau and Schönwiesner, 2018).

If both elevation and azimuth were represented in a rate code, integration could be fast and require little computational effort. In the simplest case, the normalized rates of both codes could be summed to obtain a joint representation of angular separation, in which the difference in activity elicited by a pair of sound sources is proportional to their angular separation. This unidimensional activity gradient would be insufficient to encode the joint two-dimensional direction of a sound source, but the uncertainty could be resolved by the difference in latency between azimuth and elevation processing (Fujiki et al., 2002; Bednar, Boland, and Lalor, 2017). Howver, it remains to be seen to which degree spectral cues and binaural cues are integrated in the auditory cortex, especially in more naturalistic listening situations. The adapter-probe paradigm may help to resolve this question, because it allows attenuation of populations of auditory neurons without directly affecting spatial processing.

## Acknowledgment

This preprint was created using the [LaPreprint template](https://github.com/roaldarbol/lapreprint) by Mikkel Roald-Arbøl.

